# Effects of organic fertilizer digestate on the growth of tomato (*Lycopersicon esculentum* Mill)

**DOI:** 10.1101/2022.03.30.486395

**Authors:** Branly Wilfrid Effa Effa, Dick Sadler Demikoyo, Stéphane Mibemu Guibinga, Martin Nguema Ndong, Yves Anaclet Bagafou

## Abstract

Tomatoes are known for their human health benefits. Tomato producers use chemical fertilizers to increase tomato production levels. However, the use of organic fertilizers such as digestate can achieve satisfactory yields while sustaining the soil environment. In this tomato study, the effect of three dilutions of digestate (25%, 50% and 75%) on plant height, leaf area and fruit number were compared to NPK 15 15 15. The 50% dilution of digestate had the most beneficial effect on the plants at the end of the experiment.

## 1. Introduction

Tomato (*Lycopersicon esculentum* Mill.) is one of the most popular and nutritious vegetables in the world (Falodun et Ogedegbe 2020). The nutritional and chemical composition of the tomato makes it a unique plant, it contains the following natural components vitamin C, vitamin A (as carotenoids), fiber, potassium, and the antioxidant lycopene thus the consumption of tomatoes and tomato products has a beneficial effect on human health (Burton-Freeman et Reimers 2011).

According to the FAO, the annual world production of tomatoes is estimated at about 123 million tons with a total production area of about 4.5 million ha (Gatahi 2020). While in Central Africa production was 1286716 tons with a total production area of 116804 ha in 2019 (FAOSTAT, 2021).

Vegetables need adequate growing conditions to have an optimal yield, among these conditions tomato requires adequate fertilizer for growth and yield (Singh et al. 2016). For several decades chemical fertilizers were the only ones used to increase plant yields (Uekoetter et Smil 2002), due to the negative impact of chemical fertilizers on the environment, their use should be minimized in the context of sustainable agriculture (Peyvast et al. 2008). Indeed, climate change took farmers to review their production systems, that improvement is relied on Government programs based on the Millennium Development Goals and also on Sustainable Development Goals (Tumushabe 2018). For the threat of climate change, sustainable agricultural practice has an important role to play in food insecurity (Lal 2008) despite that agricultural sector in Africa is faced with insufficient access to efficient inputs (Adenle et al. 2018). Utilization of organic fertilizer by Africans farmers could participate to sustainable agricultural, the use of organic fertilizers provides nutrients for the plant and also increase microbial activity in soil (Drinkwater et al. 1995; Angin et al. 2017). Furthermore, soil organic matter is rich in natural resources, such as nitrogen, phosphorus, and potassium (N, P, and K) in organic forms but these elements persist longer in the soil compared to their mineral (Angin et al. 2017). Thus, Ebrahimi et al. 2019 have shown that an optimized use of organic fertilizers contributes to sustainable agriculture. We used the organic fertilizer called digestate which is a by-product of methane and heat production in a biogas plant, coming from organic wastes, it could be solid or liquid depending on the biogas technology (Makádi et al. 2012).

In this study, the objective is to evaluate the effect of 3 dilutions (25%, 50% and 75%) of digestate on height, leaf area and fruits number of tomato in comparison with a chemical fertilizer of NPK 15 15 15 in the presence of no fertilizing control.

## 2. Material and Method

The study was conducted at the “*Institut de Recherches Agronomiques et Forestières”* (*0° 25’ 16, 571” N and 9° 25’ 55, 518” E*).

### Plant material

In this study we used tomato seeds of the hybrid variety MONGAL F1 (INRAE), it is high-yield variety (Ochar et al. 2019).

### Soil composition

The soil used was analyzed to determine its composition. Its composition is presented in table 1 below.

**Table 1.** Soil composition

| Composition | Quantity |
| --- | --- |
| Water content | 8.33 % |
| Total organic matter | 1.94 % |
| Total organic Carbon | 11.2 ‰ |
| Total Nitrogen | 0.28 ‰ |
| Phosphorus (P<sub>2</sub>O<sub>5</sub>) | 0.11% |
| Potassium (K<sub>2</sub>O) | 0.61 % |
| Calcium (CaO) | 1.67 % |
| Magnesium (MgO) | 0.39 % |
| Sodium (Na<sub>2</sub>O) | 1.28 % |
| Aluminium (Al<sub>2</sub>O<sub>3</sub>) | 5.15 % |
| Iron (Fe<sub>2</sub>O<sub>3</sub>) | 4.38 % |
| Manganese oxide (MnO) | 0.02 % |
| Silica (SiO<sub>2</sub>) | 61.8 ppm |
| Copper (Cu) | 22 ppm |
| Molybdenum (Mo) | 0.5 ppm |
| Zinc (Zn) | 56 ppm |
| Chloride (Cl<sup>-</sup>) | 4 g/T |
| Conductimetry/T°C | 107.3/25µS/cm |
| pH | 6.98 |
| Ratio C/N | 40 |

### Digestate composition

Analyzes carried out on the digestate in liquid form used for this study has allowed to identify elements it contains (table 2).

**Table 2.** Digestate composition

| Composition | Quantity |
| --- | --- |
| Total organic matter | 35.64 % |
| Total organic Carbon | 206.7 ‰ |
| Total Nitrogen | 7.8 ‰ |
| Phosphorus (P<sub>2</sub>O<sub>5</sub>) | 7.46% |
| Potassium (K<sub>2</sub>O) | 7.74 % |
| Calcium (CaO) | 2.38 % |
| Magnesium (MgO) | 0.86 % |
| Sodium (Na<sub>2</sub>O) | 9.62 % |
| Aluminium (Al<sub>2</sub>O<sub>3</sub>) | 0.47 % |
| Iron (Fe<sub>2</sub>O<sub>3</sub>) | 4.38 % |
| pH | 7.51 |
| Ratio C/N | 26.5 |

Three dilutions of pure digestate (table 2) were tested in our study, first dilution at 25%, second at 50% and third at 75%.

### Experimental procedures

The substrate used in this study is a mixture of soil (table 1) and sand (2V: V), sand’s grain size was between 62,5 μm and 125 μm. The substrate has been disinfected at 120°C during 20 minutes. Tomato’s seeds were sown in trays for nursery use and young plant were transplanted into plastic bags 4 weeks later and monitored for 6 weeks. Experiment was laid out in a randomized Complete Block Design (RCBD) with 8 replications and five treatments. Treatments used were a control with out fertilizer (T_0_), NPK 15 15 15 (NPK), and three dilutions of digestate in the liquid form 25% (T_1_), 50% (T_2_) and 75% (T_3_).

### Measurements of plants heights, leaves sizes and fruits diameters

Plants’ heights and leaves’ sizes were measured with a measuring tape; fruits’ diameters were measured with a caliper.

### Statistical analysis

Analysis were performed using Excel (Heiser 2005), for each parameter, the analyses compared the plants from treatment NPK with the plants from each of the treatment T_0_, T_1_, T_2_ and T_3_.

## 3. Result

In order to follow the digestate effect on plant growth, plants height, leaf area and fruits number were measured once a week in all the treatments. However, only the measurements made at the end of the experiment are presented.

### 3.1. Plant height

Plant height measurements were taken for each treatment from the first to the seventh week after transplanting. Dimension was taken from the collar to the last leaf.

Among the treatments, strong significant differences (*p-value* < 0.001) of plant height were observed between the T_2_ treatment (50% of digestate) and the NPK treatment (Figure 1) at the end of the trial (6 weeks after transplanting). Plants from treatment T_2_ having significantly larger heights than those from treatment NPK. Average value of plants height from treatments T_2_ and NPK are respectively 78.21 cm and 67.64 cm.

**Figure 1.**
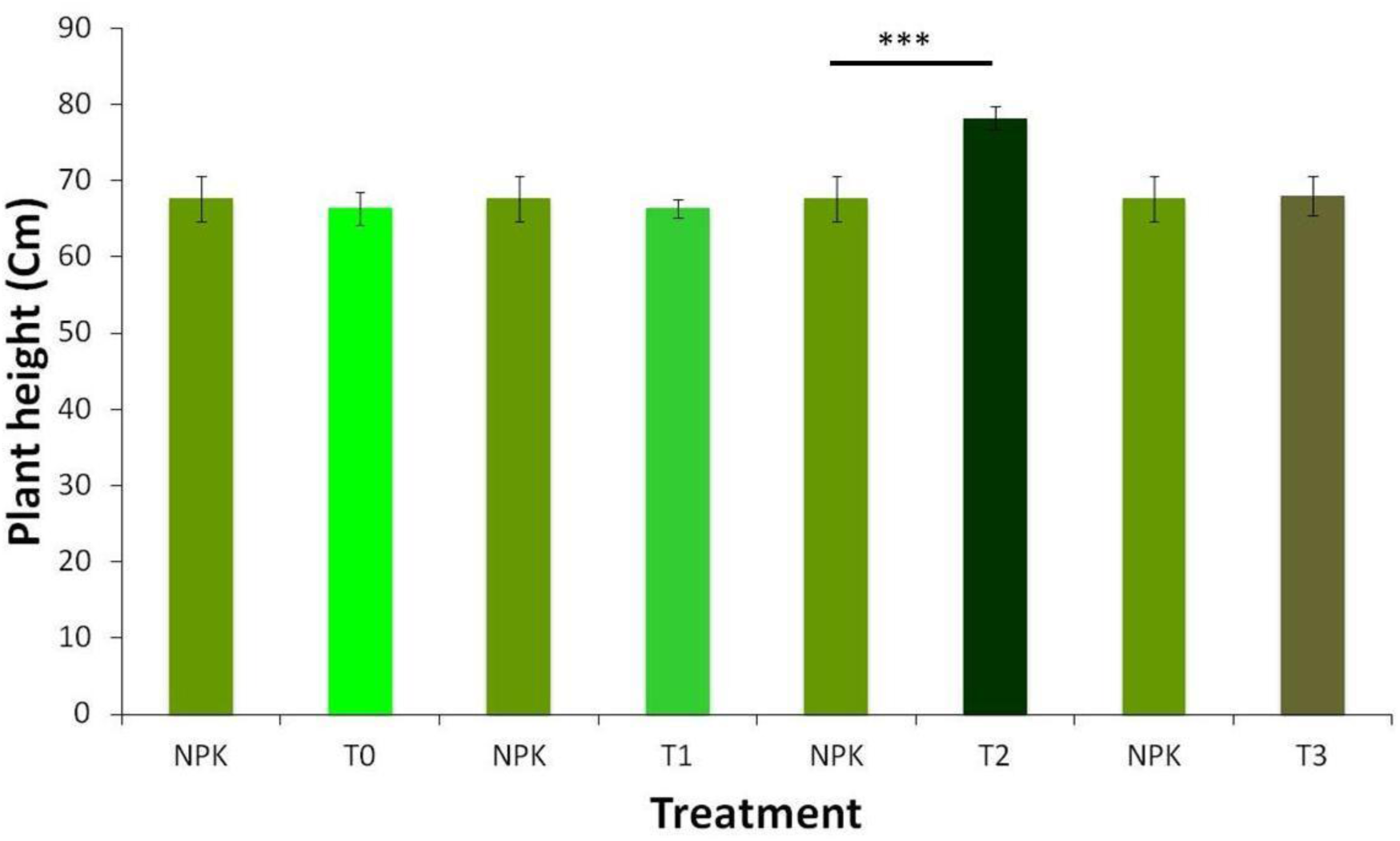
Height 6 weeks after transplantation under 5 treatments: NPK, T_0_, T_1_, T_2_, T_3_. With: NPK= fertilization with NPK 15-15-15, T_0_ = no fertilization, T_1_ = 25% of digestate dilution, T_2_ = 50% of digestate dilution, T_3_ = 75% of digestate dilution, n=8, ***: *p-value* < 0.001.

### 3.2. Leaf area

Leaf area (LA) measurements were taken on the same leaves for each treatment (Carmassi et al. 2007) the first to the seventh week after transplanting. Leaf length was measured as the distance between the insertion of the first leaflet on the rachis to the distal end whereas leaf width was measured on the widest leaflet (Carmassi et al. 2007).

Among the treatments, strong significant differences (*p-value* < 0.001) of LA were observed between T_0_ (no fertilizing) and NPK, T_1_ (25% of digestate) and NPK, also T_2_ (50% of digestate) and NPK, while significant differences (*p-value* < 0.05) of LA were observed between T_3_ (75% of digestate) and NPK at the end of the trial (Figure 2). Only plants from treatment T_2_ had a larger LA than those from NPK. Average value of plants LA from treatments T_2_ and NPK are respectively 18.73 cm^2^ and 16.86 cm^2^ at the end of the trial.

**Figure 2.**
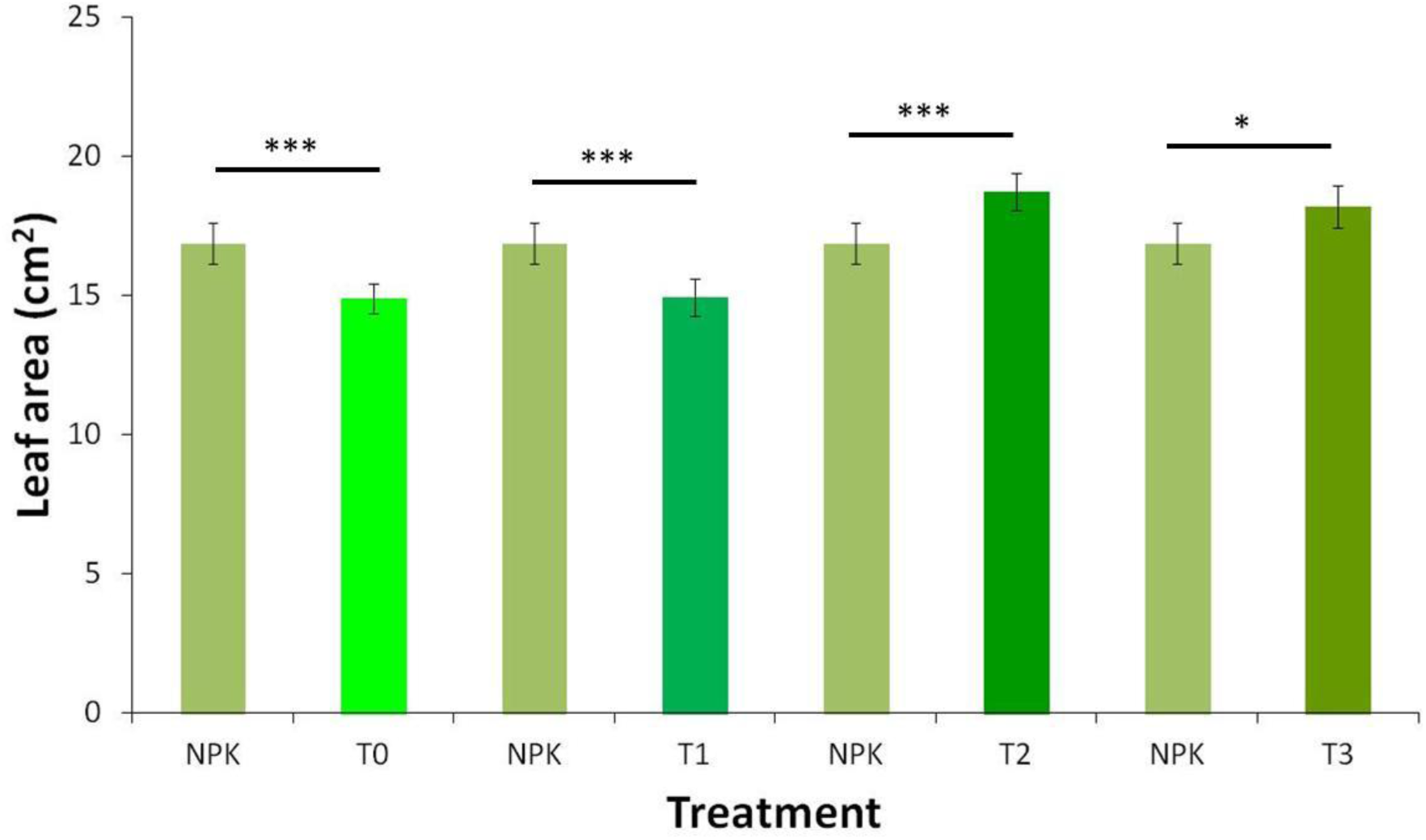
Leaf area 6 weeks after transplantation under 5 treatments: NPK, T_0_, T_1_, T_2_ and T_3_. With: NPK= fertilization with NPK 15-15-15, T_0_ = no fertilization, T_1_ = 25% of digestate dilution, T_2_ = 50% of digestate dilution, T3 = 75% of digestate dilution, n=8, ***: *p-value* < 0.001, *: *p-value* < 0.05.

### 3.3. Fruit diameter

Fruit diameter measurements were taken on all fruits for each treatment at the end of the trial.

Among the treatments, strong significant differences (*p-value* < 0.01) of fruit diameter were observed between T_0_ and NPK, T_1_ and NPK, and also significant differences (*p-value* < 0.05) between T_3_ and NPK. Despite these differences, the plants from the NPK treatment having the largest diameters (figure 3). However no difference of fruit diameter was observed between T_2_ and NPK at the end of the trial (figure 3).

**Figure 3.**
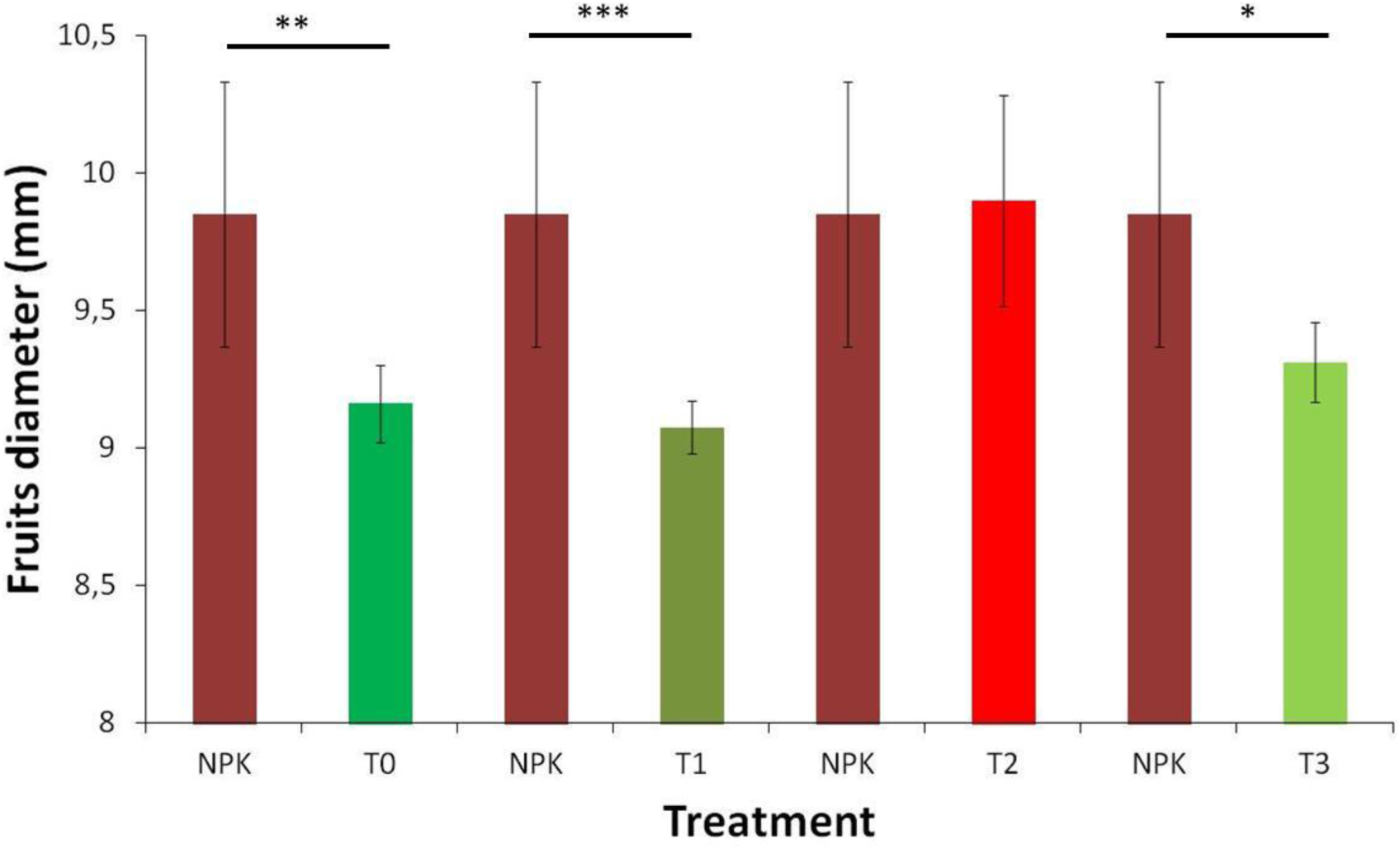
Fruits diameter 6 weeks after transplantation under 5 treatments: NPK, T_0_, T_1_, T_2_ and T_3_. With: NPK= fertilization with NPK 15-15-15, T0 = no fertilization, T_1_ = 25% of digestate dilution, T_2_ = 50% of digestate dilution, T_3_ = 75% of digestate dilution, n=8, ***: *p-value* < 0.001, **: *p-value* < 0.01, *: *p-value* < 0.05.

**Figure 4.**
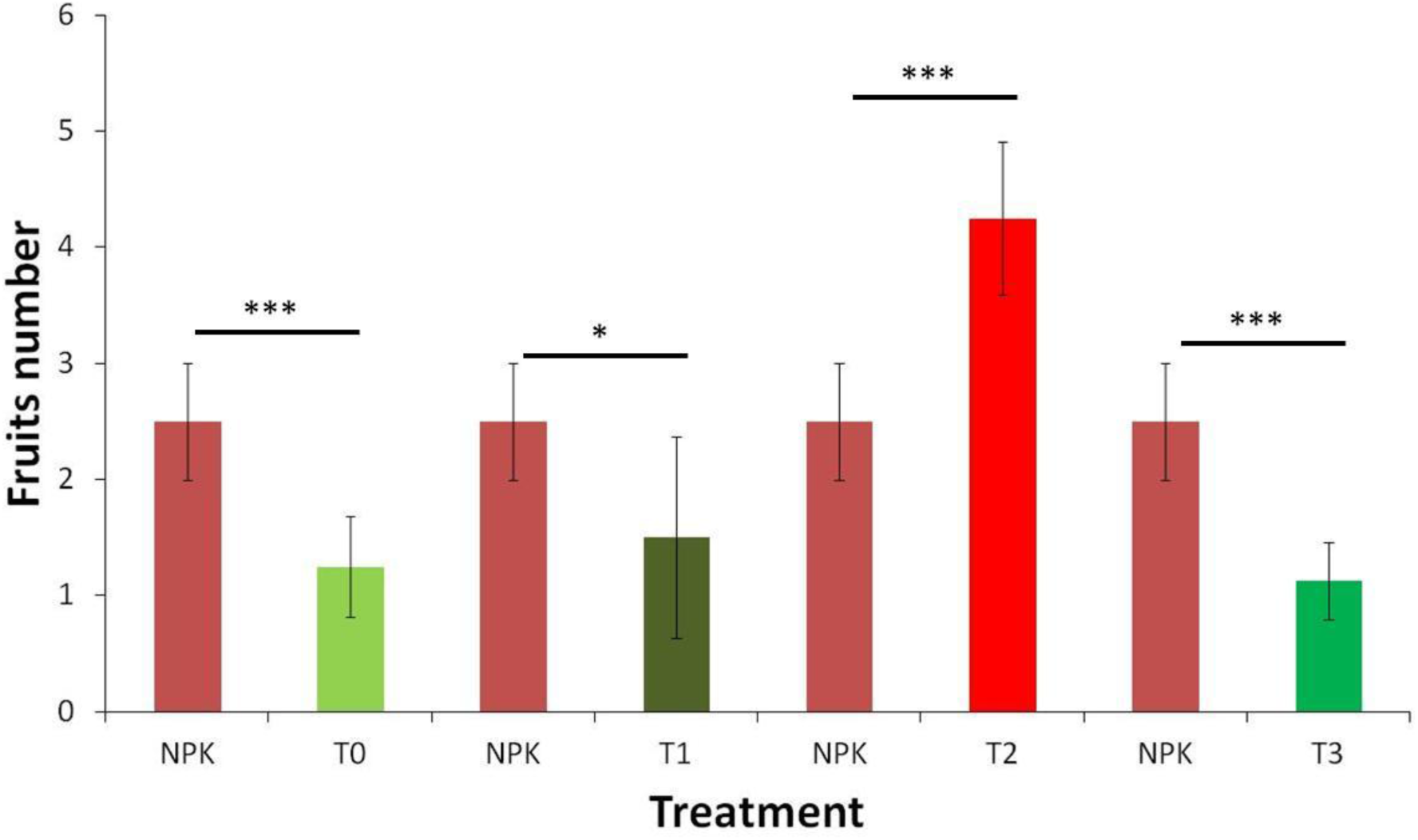
Fruits number 6 weeks after transplantation under 5 treatments NPK, T_0_, T_1_, T_2_ and T_3_. With: NPK= fertilization with NPK 15-15-15, T_0_ = no fertilization, T_1_ = 25% of digestate dilution, T_2_ = 50% of digestate dilution, T_3_ = 75% of digestate dilution, n=8, ***: *p-value* < 0.001, *: *p-value* < 0.05.

### 3.4. Fruit number

Fruit number (FN) was counted for each treatment at the end of the trial.

Among the treatments, strong significant differences (*p-value* < 0.001) of fruit diameter were observed between T_0_ and NPK, T_2_ and NPK, T_3_ and NPK, and also significant differences (*p-value* < 0.05) between T_1_ and NPK. Only plants from treatment T_2_ had a larger FN than those from NPK. Average value of plants FN from treatments T_2_ and NPK are respectively 4.25 and 2.5 at the end of the trial.

## 4. Discussion

There is often variability between soils in different regions and even within a field (Shuman et al. 1980). In view of the high population growth in Africa, the use of optimal soil management methods would help to increase the yields of agricultural production (Lal 2008). Utilization of organic fertilizer by Africans farmers could participate to sustainable agricultural, the use of organic fertilizers provides nutrients for the plant and also increase microbial activity in soil (Drinkwater et al. 1995; Angin et al. 2017).

In this study, it was observed that the plants from the T_2_ (50 % of digestate) treatment had the best vegetative development and the highest yield at the end of the trial compared to NPK (NPK 15 15 15), T_0_ (25 % of digestate) and T_3_ (75 % of digestate). This result is in agreement with the work of Agbede et al. 2019 who obtained better vegetative development and yield with green manure used as organic fertilizer compared to the NPK 15 15 15 treatment. Digestate could therefore be used as a substitute for chemical fertilizers insofar as plants grown in soil fertilized with digestate have significantly higher yields than plants grown in soil fertilized with NPK. Furthermore, its use preserves the soil biomass, unlike chemical fertilizers (Angin et al. 2017).

However an optimal concentration of the digestate would result in good quality plants and high yields. In our trial, T_2_ would be the optimal treatment in view of the results obtained with plants from treatments T_0_ (no fertilizing), T_1_ (25% of digestate) and T_3_ (75 % of digestate).

## 5. Conclusion

Our study showed that the application of digestate (organic fertilizer) as a fertilizer at an optimal dose during tomato cultivation results in higher yields than those obtained with NPK 15 15 15. The digestate could therefore replace NPK (chemical fertilizer) which has a negative impact on the environment.

## Acknowledgements

Our acknowledgement goes to the **General Direction of Livestock** for the provision of digestate and the colleagues of **IRAF** who contributed to the realization of this work.

